# Vesicular delivery of the antifungal antibiotics of *Lysobacter enzymogenes* C3

**DOI:** 10.1101/344838

**Authors:** Paul R. Meers, Carol Liu, Rensa Chen, William Bartos, Julianne Davis, Nicole Dziedzic, Jason Orciuolo, Szymon Kutyla, Maria Jose Pozo, Deepti Mithrananda, Dominick Panzera, Shu Wang

## Abstract

*Lysobacter enzymogenes* C3 is a predatory strain of gram-negative gliding bacteria that produces antifungal antibiotics by the polyketide synthetic pathway. Outer membrane vesicles (OMV) are formed as a stress response and can deliver virulence factors to host cells. The production of OMV by C3 and their role in antifungal activity are reported here. Vesicles in the range of 130-150 nm in diameter were discovered in the cell-free supernatants of C3 cultures. These OMV contain molecules characteristic of bacterial outer membranes, such as lipopolysaccharide and phospholipids. In addition, they contain chitinase activity and essentially all of the heat stable antifungal activity in cell supernatants. We show here that C3 OMV can directly inhibit growth of the yeast *Saccharomyces cerevisiae* as well as the filamentous fungus *Fusarium subglutinans*. The activity is dependent on physical contact between OMV and the cells. Furthermore, fluorescent lipid labeling of C3 OMV demonstrated transfer of the membrane-associated probe to yeast cells, suggesting the existence of a mechanism of delivery for membrane-associated molecules. Mass spectrometric analysis of C3 OMV extracts indicates the presence of molecules with molecular weights identical to some of the previously identified antifungal products of C3. These data together suggest that OMV act as an important remote mobile component of predation by *Lysobacter*.

**Importance:** The data presented here suggest a newly discovered function of outer membrane vesicles (OMV) that are produced from the outer membrane of the bacterial species *Lysobacter enzymogenes* C3. We show that these OMV can be released from the surface of the cells to deliver antibiotics to target fungal organisms as a mechanism of killing or growth inhibition. Understanding the role of OMV in antibiotic delivery can generally lead to improved strategies for dealing with antibiotic-resistant organisms. These results also add to the evidence that some bacterially produced antibiotics can be discovered and purified using methods designed for isolation of nanoscale vesicles. Information on these systems can lead to better identification of active molecules or design of delivery vehicles for these molecules.

## Introduction

*Lysobacter enzymogenes* C3 (referred to here as C3) was originally identified as *Stenotrophomonas maltophilia* (1), and broad-range antifungal activity has been associated with its predatory lifestyle. Because this activity acts against pathogenic fungi that attack various crop species, C3 has been under development as a biocontrol organism to combat several plant fungal diseases (2,3). This antifungal activity may also involve molecules that can act as therapeutic agents for human fungal diseases (4). C3 is known to produce bioactive molecules through the polyketide synthase /non-ribosomal peptide synthetase (PKS/NRPS) pathway (3). Some of these molecules, particularly polycyclic tetramate macrolactams, are stable to heat, can be obtained by extraction with organic solvents, and have been demonstrated to kill or inhibit the growth of various specific fungi. This activity was originally identified as a complex that could be purified by ammonium sulfate precipitation, and was designated heat stable antifungal factor (HSAF) (3). Dihydromaltophilin (3,5) had been initially identified as one of the major C3 antifungal antibiotics in HSAF, and is now essentially synonymous with HSAF. In addition, other polycyclic tetramate macrolactam products of C3, such as alteramide A and B, have been identified with apparently different antifungal specificities and mechanisms of activity (6). The process by which HSAF is delivered to fungal cells has not been elucidated, although it has been suggested that one or more of the well-defined type I-IV secretory complexes may mediate secretion or injection into host cells (7).

Gram-negative bacteria are also known to produce outer membrane vesicles (OMV) under certain circumstances. These vesicles bud from the outer membrane, often as a stress response (8-11), and can contain a number of different components, including toxins (12), signaling molecules (13), genetic material (14-16), and self-defense compounds, such as enzymes that can degrade antibiotics (17). Accumulating evidence indicates that OMV play a major role in host-pathogen interactions (18, 19). In some species, OMV harbor small-molecule anti-host antibiotics (20, 21), while in others the anti-host activities are primarily enzymatic (9, 22). In particular, vesicle production has been previously documented in *L. enzymogenes sp.* XL1, showing the association of host-lytic enzymes with the vesicles (23, 24). However, the production of OMV by the C3 biocontrol strain has not been previously investigated.

We report here that C3 grown in liquid culture forms what appear to be outer membrane vesicles (OMV) that contain most if not all of the heat-stable antifungal activity associated with supernatants from C3 cells. C3 OMV were found to be approximately 130-150 nm in diameter, contain phospholipids and lipopolysaccharide, DNA, and a collection of associated proteins, including a chitinase activity. Heat-stable antifungal activity against a filamentous fungus and yeast were observed, and experiments exploring the mechanism of transfer to these host cells are presented. Purified OMV extracts yielded molecules with mass spectra expected for known C3 PKS/NRPS pathway products with antifungal activity. These data demonstrate vesicular packaging of intrinsically synthesized small molecule antibiotics by this predatory species.

## RESULTS

### The C3 strain produces outer membrane vesicles

The presence of OMV in the cell-free supernatants of C3 was tested under several conditions. Putative OMV could be obtained either by directly sedimenting cell-free supernatant to obtain an OMV pellet or by ultrafiltration of the supernatants through a 300 kDa cutoff filter (equivalent to a spherical protein particle cutoff of approximately 7-8 nm). In the latter case, the material collected from the filter was then sedimented and washed for further analysis. Additionally, minimal medium (MM) or 10% tryptic soy broth (TSB) was found to support C3 OMV production, whereas a rich medium such as Luria broth (LB) did not. Vesicle material could only be obtained in late log phase or stationary phase culture and not in mid-log phase.

Observation of the washed, resuspended material by light microscopy revealed particles, apparently less than 1 micron (Figure 1). It was possible to label these particles with a well-characterized probe, diIC18 (see methods), which resulted in labeled submicron particles that could be observed by fluorescence microscopy (Figure 1). Importantly, an enhancement in fluorescence of the probe was observed (approximately 5-10-fold, data not shown), consistent with the behavior expected for insertion into a hydrophobic membrane environment. These data suggested that the observed particles were indeed vesicles with a delimiting membrane, consistent with OMV. Samples were also analyzed by negative stain transmission electron microscopy (EM), showing apparently aggregated particles/vesicles with diameters in the 100-150 nm range (Figure 1). Aggregation, fusion and/or other distortions of the negatively charged vesicles may be an artifact of the negative stain EM preparation, perhaps as a result of the presence of polyvalent positive uranyl ions in the procedure.

**Figure 1:**
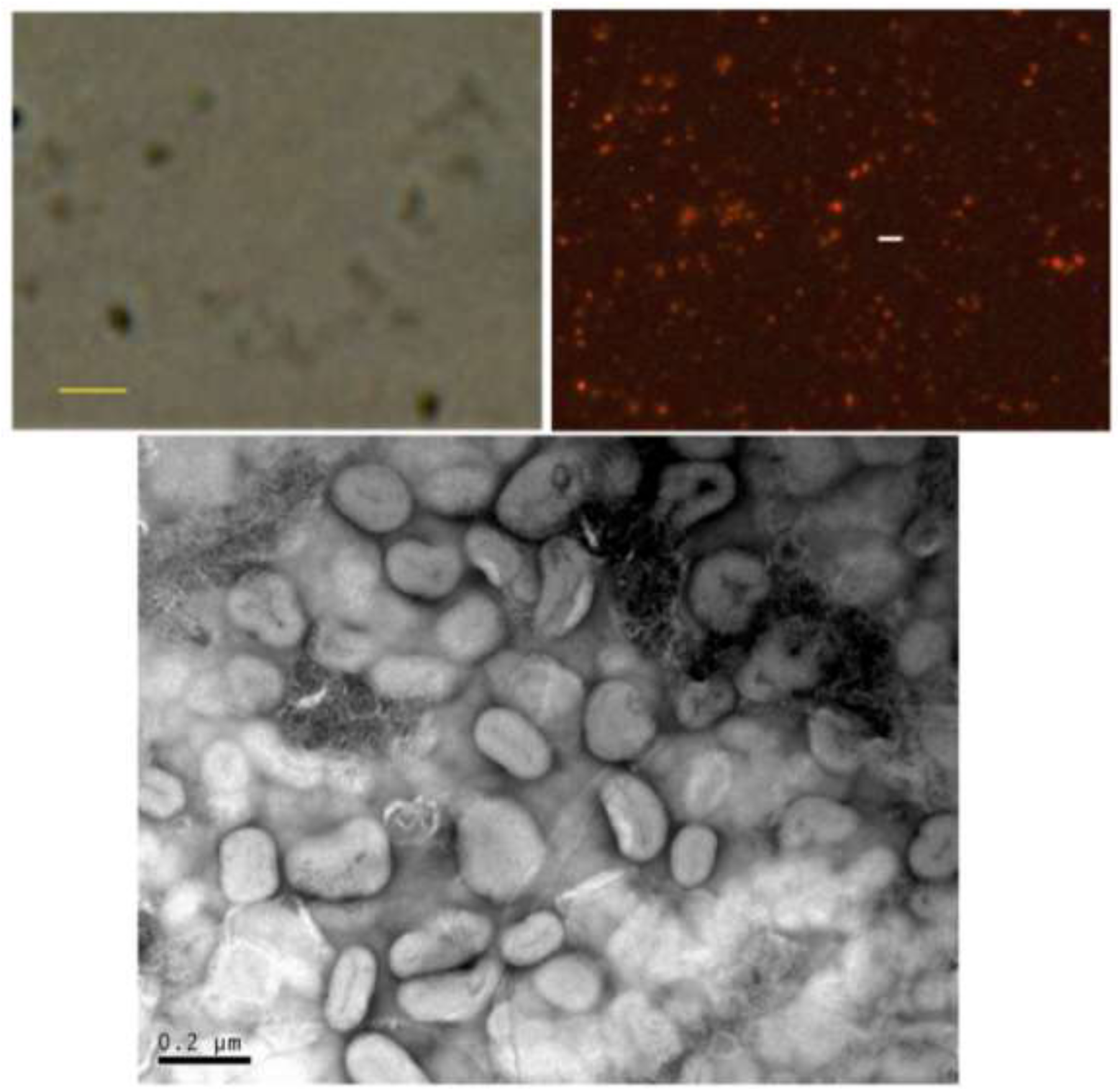
Microscopic images of vesicular material from C3 cell culture supernatants. Top left: 800x phase contrast image of unlabeled vesicles. Bar represents 1 micron. Top right: Fluorescence photomicrograph (800x) of vesicles labeled with diIC18. Bar represents 2 microns. Bottom panel: Negative stain electron microscopic images of C3 OMV preparation. Bar represents 0.2 microns.

Additionally, OMV from washed pellets were sedimented on a discontinuous iodixanol gradient (equal volumes of 15, 25, 40 and 54%) and migrated into a visible band at the interface between 15 and 25% iodixanol (approximately 1.18 and1.3 g/cm^3^, respectively, data not shown). This density is consistent with the buoyant densities of previously identified OMV of other bacteria (25,26).

Size and charge of the particulate samples were also analyzed by dynamic light scattering and electrophoretic mobility, respectively. The data for three separate preparations revealed a homogeneous and narrow size distribution with a z-average size of approximately 135 nm, a low polydispersity index (PDI) of approximately 0.09 and a notable lack of particles of other sizes (Figure 2). Zeta potential measurements indicated a negative surface potential of approximately - 10 mV.

**Figure 2:**
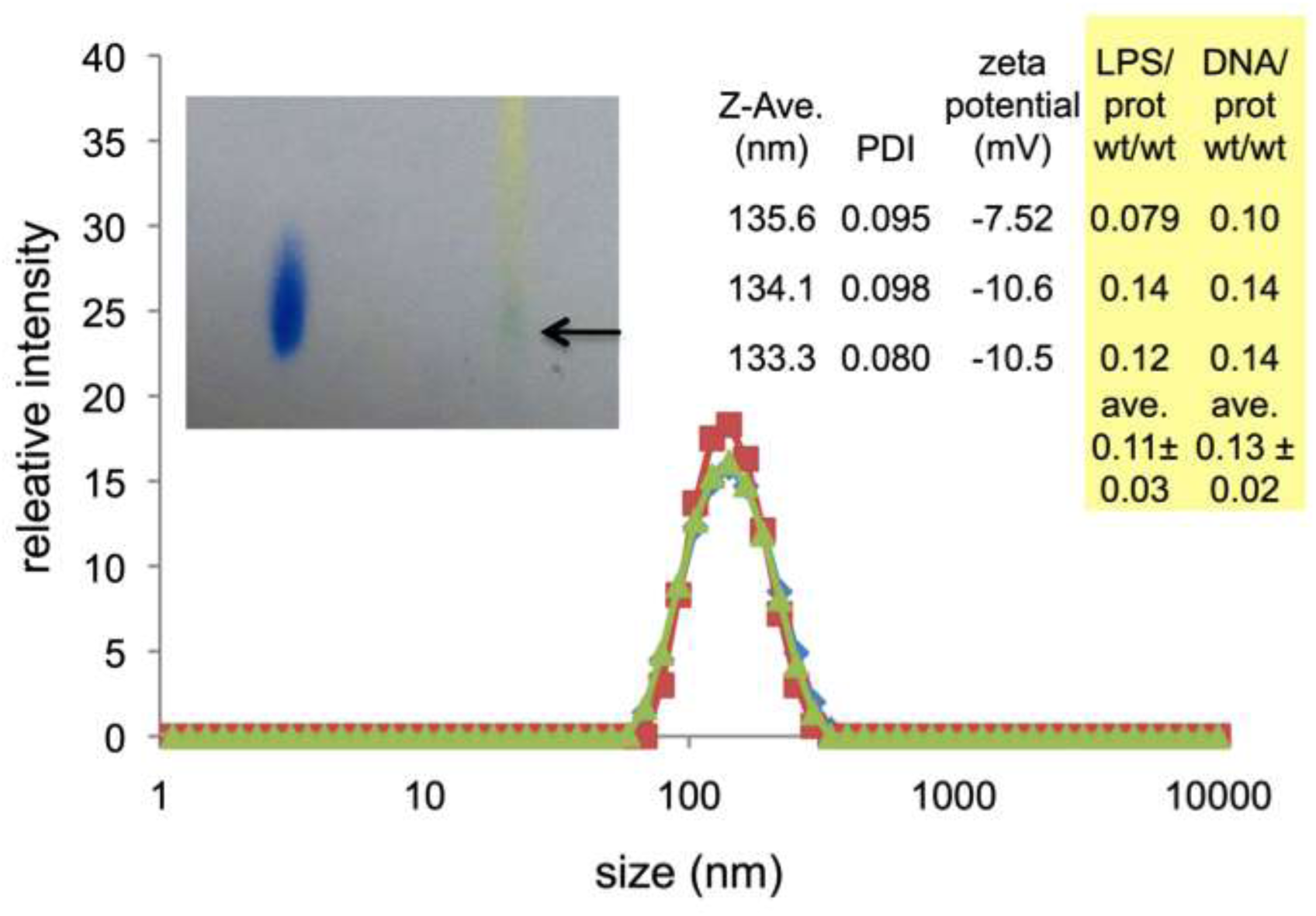
Size, charge and composition of three OMV preparations. Graph represents the distribution of vesicle sizes detected by dynamic light scattering with a peak in z-average intensity at approximately 135 nm. Inset table at the right shows z-average size for each sample, polydispersity index (PDI), as well as zeta potential. Data for estimated LPS/protein and DNA/protein ratios are also shown for each preparation. Left inset shows evidence for phospholipids from a Bligh Dyer extract of C3 OMV as detected with thin layer chromatography with Zinzadze reagent staining. Left sample: standard POPG (1-palmitoyl-2-oleoyl-sn-glycero-3-phospho-(1′-rac-glycerol)), Rf ~ 0.41. Right sample: Bligh Dyer extract of C3 OMV where arrow denotes blue phosphate positive spot, Rf ~ 0.42.

Outer membrane vesicles of bacteria would also be expected to contain lipids characteristic of the cell envelope. C3, like most gram-negative bacteria, displays lipopolysaccharide (LPS) on its surface. Therefore, the presence of LPS in these vesicles and its relationship to the protein content was tested in the same three preparations (see Figure 2). For spherical vesicles of approximately 135 nm in diameter, the expected LPS/protein ratio based strictly on geometrical considerations would fall in the range of 0.08 to 0.66 (see Methods), depending on whether any protein is encapsulated in the interior of the vesicle, as well as the membrane protein content. These preparations of C3 OMV contained an estimated average ratio of 0.11±0.03 LPS/protein g/g based on an *E. coli* LPS standard and the Bradford protein assay. These data are consistent with a considerable amount of protein, both in the membrane and encapsulated inside the vesicles (see calculations in Materials and Methods). Additionally, Bligh Dyer extracts (27) of OMV preparations that were analyzed by thin layer chromatography appeared to show components consistent with phospholipids commonly found in bacterial outer membranes, such as phosphatidylglycerol or phosphatidylethanolamine (Figure 2 inset). The presence of outer membrane lipids further corroborates the identity of the isolated material as OMV.

Protein content of the vesicles was analyzed by polyacrylamide gel electrophoresis, western blotting immunoassay and by an assay for chitinase activity. Gel electrophoresis showed major protein bands as documented in Table 1 (data not shown). Chitinase activity was consistently detected in washed OMV preparations from chitin-induced cultures as monitored by a fluorogenic substrate assay (see Materials and Methods). Treatment of OMV with proteinase K did not entirely eliminate chitinase activity, indicating that at least some portion of the chitinase may be encapsulated inside the OMV (Table 1). 10X freeze-thaw and washing of these OMV by repeated sedimentation also resulted in retrieval of approximated 20% of the total OMV-associated chitinase activity. These data suggest the possibility of a membrane-associated form of chitinase.

**Table 1:** Preliminary characterization of components of C3 OMV. Experimental details are given in Materials and Methods.

| OMV<br>component | content identification | entrapped material |
| --- | --- | --- |
| protein | Major bands: 28, 31, 37,<br>53, 70 kDa<br>chitinase activity<br>observed | ~80-89 % chitinase activity<br>protected from proteinase K<br>digestion |
| nucleic acids | 0.13 g DNA/g protein | ~ 100 % inside (not accessible<br>to DNase) |
| lipids | 0.11 g LPS/g protein<br>phospholipids present |  |

OMV preparations were also tested by western blotting for the presence of pilA, the major component of *Lysobacter* pili. No evidence for the existence of pilA was found in OMV preparations, while a strong band could be seen in an equal protein concentration of whole cell preparations (Figure 3). These data confirm that C3 OMV are truly a result of the budding of specific portions of the outer membrane, and not just outer membrane fragments, which may retain pili.

**Figure 3:**
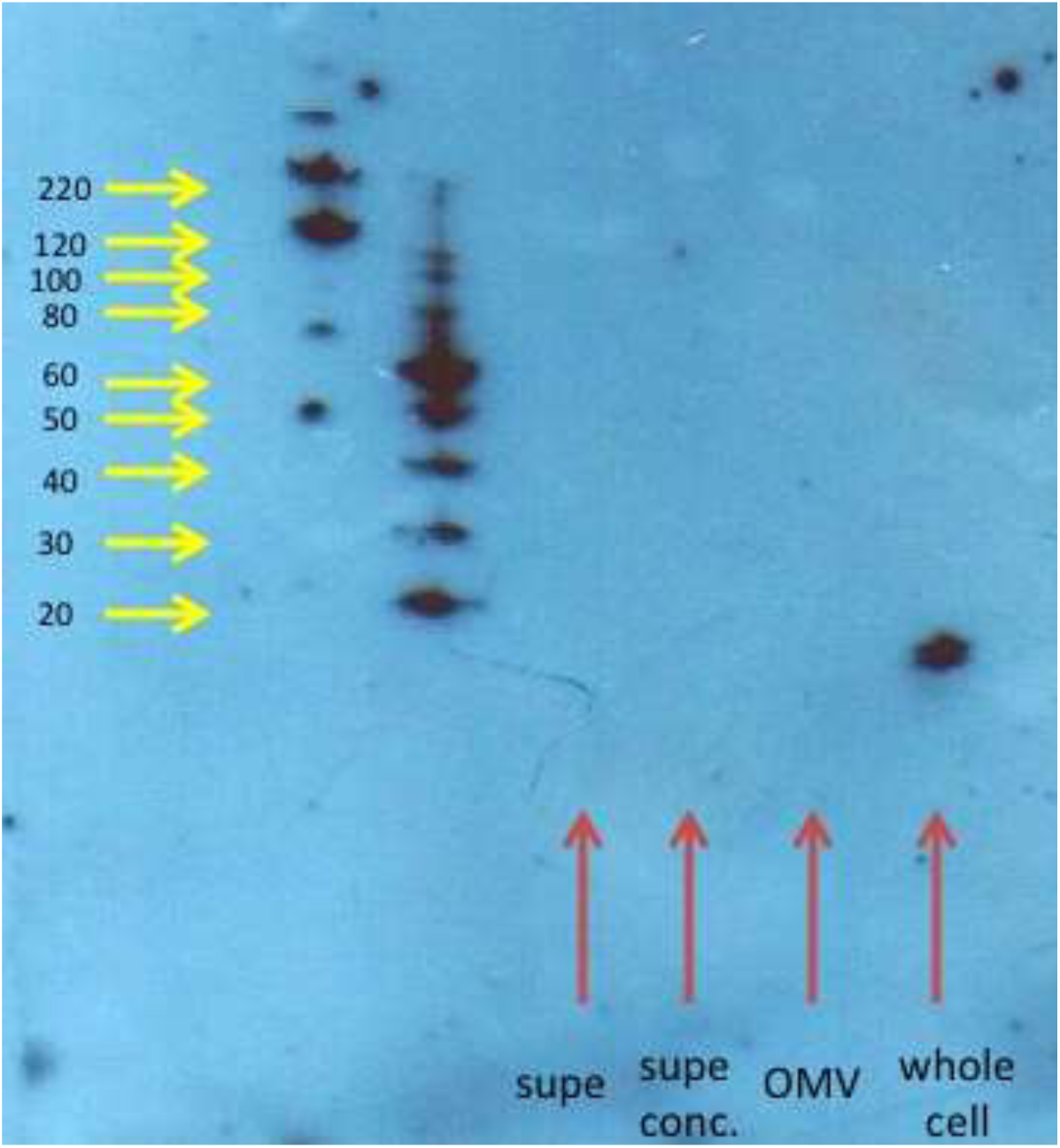
Western blot analysis of C3 OMV, whole C3 cells, and supernatants from C3 OMV. An 8-25% gradient acrylamide gel was used to separate samples, and the blot probed with an antibody against a peptide within the pilA protein. The expected molar mass of pilA is 14 kDa. Molecular weights of standards are shown with yellow arrows. Samples are shown with red arrows. Equal amounts of protein from OMV and whole cells were loaded on the gel.

Nucleic acids are also commonly found to be associated with OMV both externally and internally (14-16, 25). DNA was detected in C3 OMV preparations by standard ethanol precipitation procedures, and the presence of DNA was also observed via a fluorogenic DNA-binding probe (see Material and Methods and tabular portion of Figure 2). When intact OMV were treated with DNase I, most of the DNA in the OMV preparations was inaccessible to the enzyme, indicating encapsulation of most DNA in the interior of the vesicles. Although the presence of a small amount of RNA cannot be ruled out, treatment of nucleic acids isolated from OMV with RNase-free DNase did not yield detectable material.

In summary, a particulate fraction with a well-defined z-average diameter of approximately 135 nm is obtained from C3 culture supernatants under limiting nutrient conditions. The existence of membrane lipids, enhancement of membrane probe fluorescence, observation of encapsulated material and other components common to OMV of gram negative bacteria, strongly suggest that OMV are a common product of *L. enzymogenes* C3.

### Antifungal activity is associated with OMV

A yeast growth assay (*Saccharomyces cerevisiae*, strain S30) was used as the primary test for antifungal activity of fractions prepared from C3 culture. Activity against *Fusarium subglutinans* was also tested in some cases (see below). Initially, extraction methods similar to those previously reported to retrieve HSAF from cell supernatants were used (28). Specifically, ethyl acetate extracts of the cell-free supernatants were obtained, dried and redissolved in a smaller volume of ethanol and tested in yeast growth assays as described in Materials and Methods. Antifungal activity was observed from late log or stationary phase C3 supernatants, while no activity (or OMV) was found in mid-log phase supernatants (data not shown).

Cell-free supernatants were also fractionated by ultrafiltration through a 300 kDa cutoff filter. This resulted in an observable film of material on the filter surface. When the filter material was resuspended in an equal volume of medium as compared to the filtrate and extracted with ethyl acetate as above, the antifungal activity assay showed that >90% of the activity was recoverable from the surface of the filter (Figure 4). The activity present in resuspended OMV material could be further sedimented by high-speed centrifugation (140,000 × g). The washed pellets containing OMV were then also analyzed for protein content and activity. These data also showed that all detectable activity was associated with the OMV (see below).

**Figure 4:**
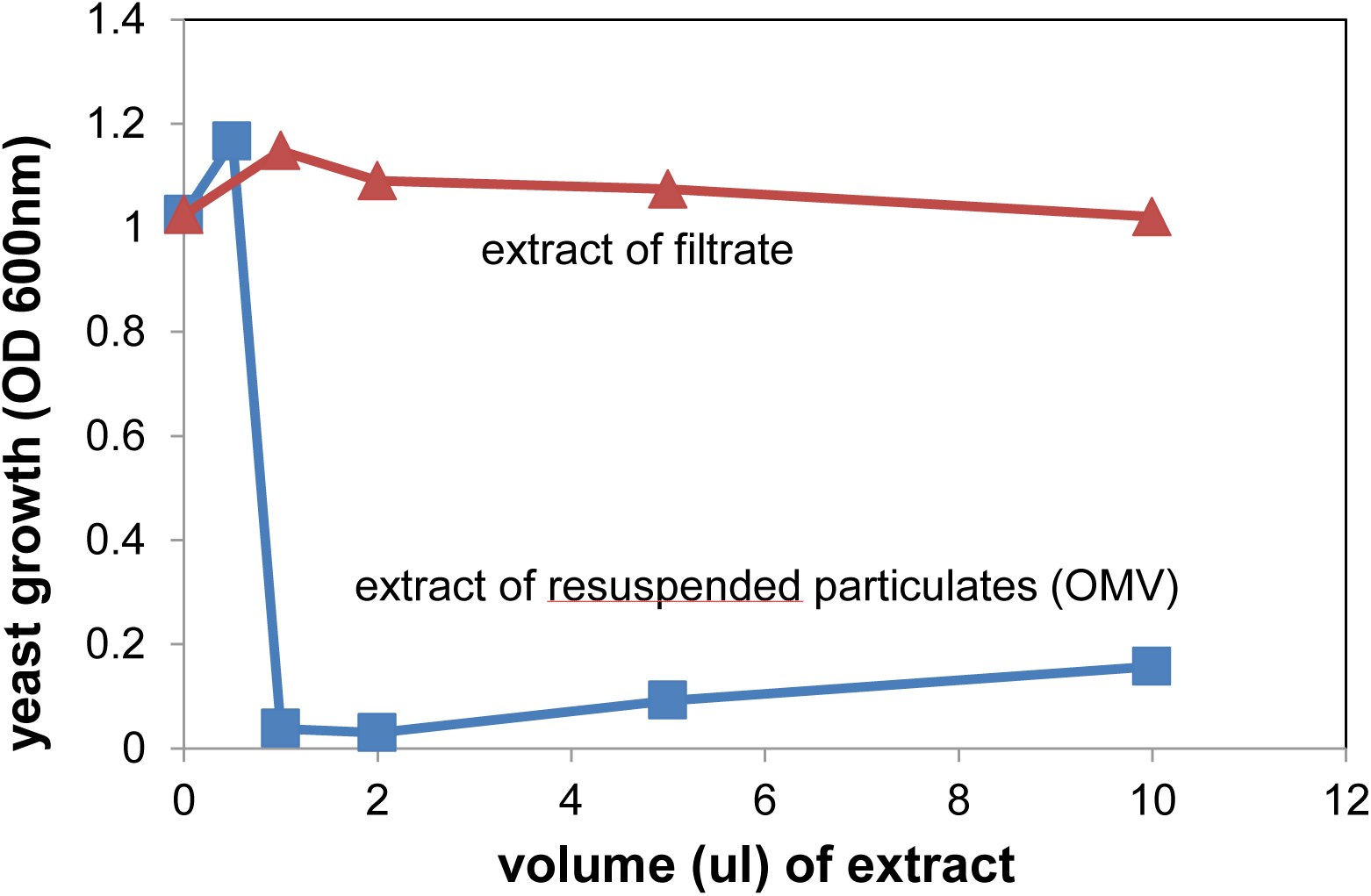
Yeast growth inhibitory activity of extract samples from filtrates and resupended OMV. Resuspended OMV and filtrate fractions (300 kDa cutoff) were extracted under identical conditions, dried and redissolved in equal volumes of ethanol. Ethanol aliquots were added to dilute yeast cultures (S30 strain) in the volumes indicated, and overnight growth was monitored by turbidity at 600 nm. More details are given in Materials and Methods.

The nature of the antifungal activity in these OMV pellets was further investigated. Figure 5 shows the dose response in terms of OMV protein concentration. We found that active OMV could be isolated from MM as well as 10% TSB, but in MM, addition of colloidal chitin to the culture medium was necessary to obtain OMV with maximal specific activity. OMV from chitin-induced cultures also showed chitinase activity. When the OMV samples were heat-treated at 70 °C for 30 minutes, all of the yeast growth-inhibitory activity remained intact (Figure 5) while all chitinase activity was destroyed (not shown). Therefore it can be concluded that the observed antifungal activity is solely due to heat-stable factors that are likely to be small molecules, at least under the conditions of these experiments. Additionally, OMV were isolated from a mutant strain of C3 in which the PKS/NRPS pathway is disrupted (designated HSAF^-^).

**Figure 5:**
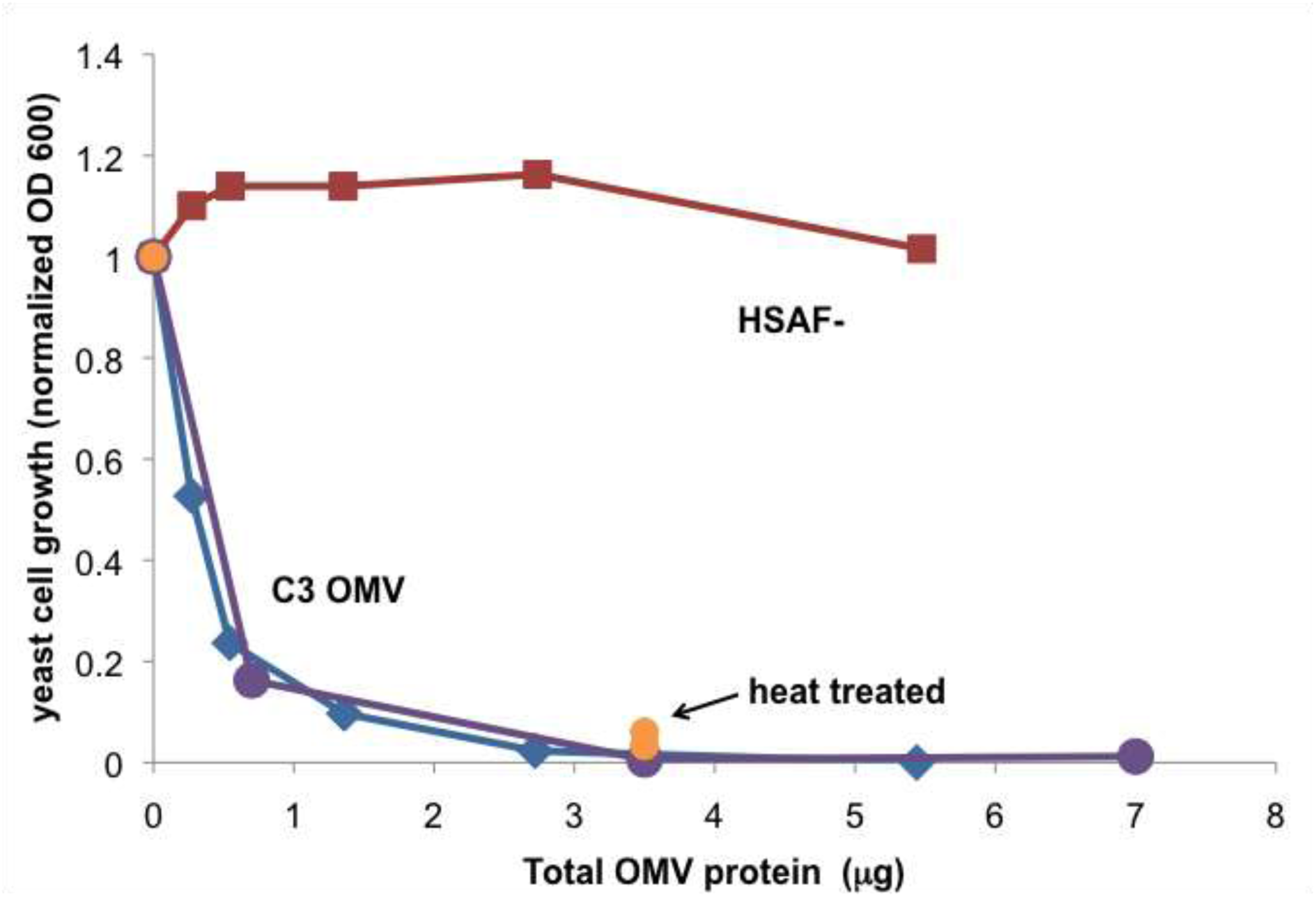
Specific activity of OMV preparations in yeast growth assays. Two separate preparations of C3 OMV were isolated from culture supernatants (blue and violet, “C3 OMV”). Orange data points represent OMV that were heat treated at 70 °C before testing for activity. OMV isolated from C3 HSAF^-^ mutant cells (HSAF^-^ OMV, red). No activity was observed in samples of similar volume from the OMV-free supernatants of these preparations. Data are representative of many preparations of OMV.

These OMV showed no antifungal activity. These data taken together strongly suggested that C3 OMV package heat stable antifungal molecules, such as dihydromaltophilin, and that the observed antifungal activity is solely due to these molecules.

Further characterization and confirmation of OMV-association of antifungal activity was performed using isopycnic sedimentation on iodixanol density medium. Based on the observation of the density behavior of OMV samples mentioned above, they were loaded onto pure 25% iodixanol to test whether more dense antifungal protein particles may have been mixed with the relatively light vesicles. In this case, a band of OMV material could be visually observed near the top of the 25% iodixanol solution after high-speed centrifugation. When pre-labeled with diIC18, the fluorescence of labeled OMV also migrated to a position near the top of the 25% iodixanol (not shown). Correspondingly, yeast antifungal activity also remained near the top of the 25% iodixanol, as did a large portion of the protein (Figure 6). Therefore, it would appear that the C3 antifungal activity is exclusively associated with light fractions, again consistent with OMV localization.

**Figure 6:**
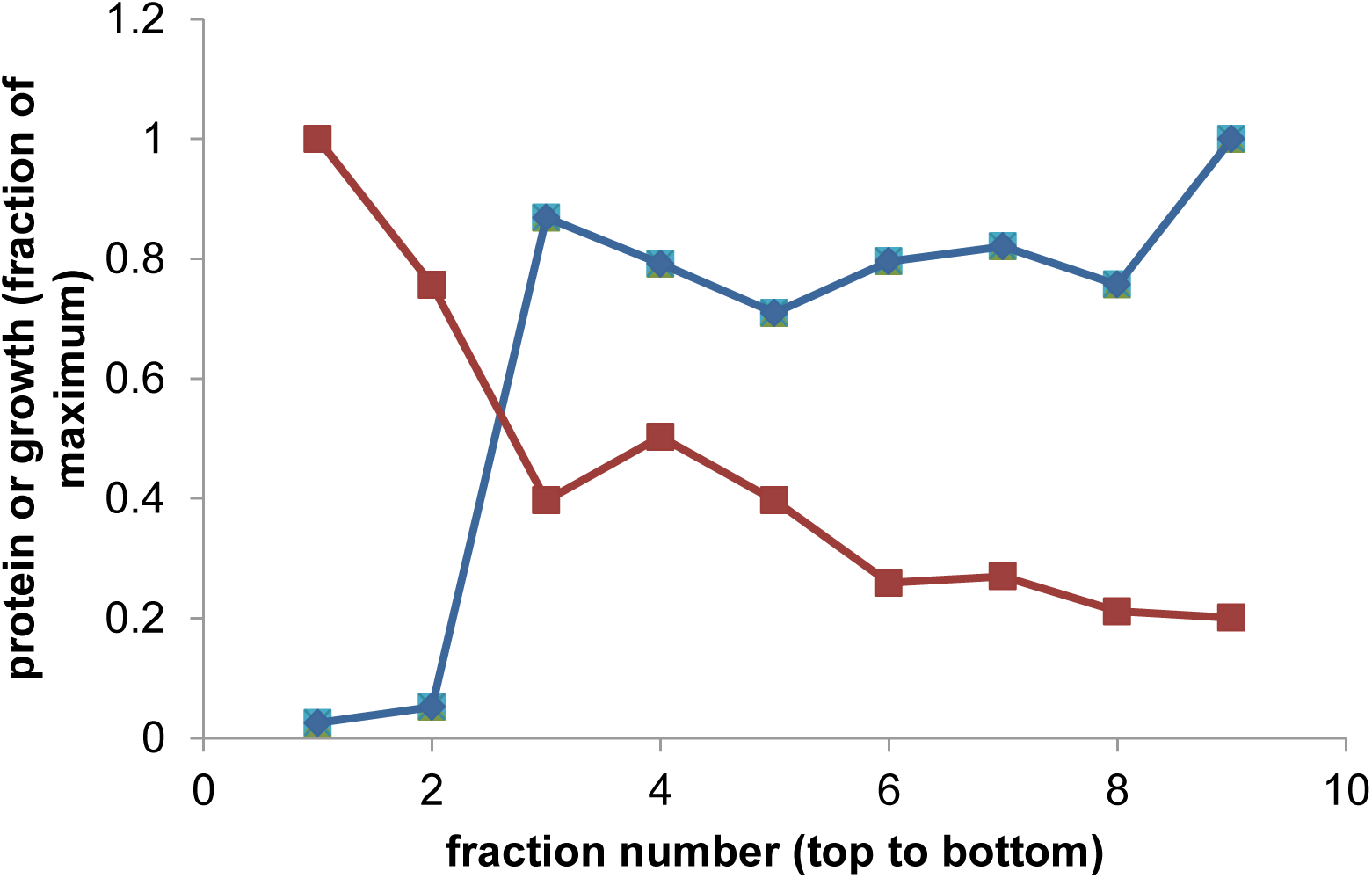
Density-based fractionation of antifungal activity and protein concentration of OMV samples. OMV samples loaded on 25% iodixanol were recovered after centrifugation in the fractions shown, where 1-10 run from the top to the bottom of the centrifuge tube, corresponding to least to most dense material. The protein concentration and effect of each fraction on yeast cell growth was determined and plotted on a fraction of maximum scale.

C3 is known to exert antibiotic activity against filamentous fungi. Therefore OMV preparations were also tested against a filamentous fungus, *Fusarium subglutinans*. This species was grown on agar plates and the activities of OMV preparations were tested by placing the samples in wells in the agar. As can be seen by the lack of mycelia around the OMV well, *Fusarium* growth was also inhibited by the C3 OMV (Figure 7). Although, the format of the assay is quite different, the OMV protein concentration required to inhibit *Fusarium* growth was generally in a range similar to the concentrations needed to inhibit yeast growth.

**Figure 7:**
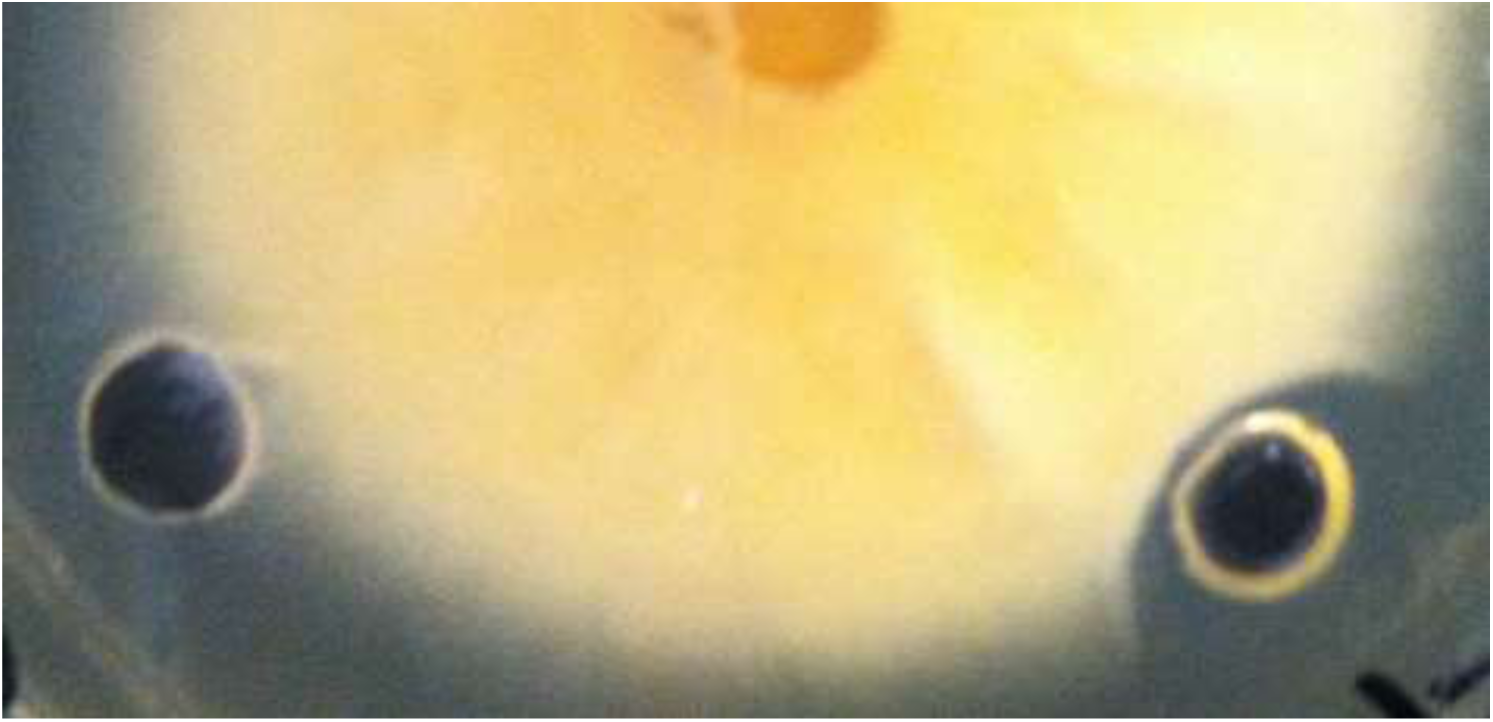
Growth inhibition of mycelia of *Fusarium subglutinans* by OMV. *Fusarium* was grown from the center of an agar plate and wells in the agar containing control buffer (left) or C3 OMV (right) were evaluated. Clear zone around the right-hand well indicates growth inhibition.

### OMV-mediated delivery of antifungal compounds to yeast cells

A corollary of OMV localization of antifungal activity is the hypothesis that there is a mechanism by which OMV can deliver the relevant compounds to yeast and other fungal cells. The requirements for delivery would depend critically on how antifungal antibiotics are associated with the OMV. The effect of freeze-thawing on the OMV activity was used as an initial crude test of localization. If activity is localized in a water-soluble form in the interior of the vesicles, freeze-thawing should destabilize the membranes and release the encapsulated molecules or complexes. 10x freeze-thawing and washing of the OMV did not affect the ability of OMV to inhibit yeast growth (data not shown), suggesting that the active molecules may be membrane-bound.

If activity is membrane-bound and not easily diffusible through aqueous media, delivery may require close contact between OMV and host cells. A filter-based activity assay was used to test this concept. Yeast cells and C3 OMV were placed on opposite sides of a 100 kDa filter (“trans-filter”) or the same side (“cis-filter”), and the ability of the yeast cells to grow was measured. Short (5.5 hr.) and long (overnight) incubation times were used. At either incubation time, the 100 kDa filter did not allow diffusion of active molecules from OMV to reach the yeast cells so as to inhibit their grow, resulting in a relatively high turbidity of the trans-filter yeast culture (OD600 = 0.176±0.037; 0.228±0.0007, respectively). Under the same conditions, incubation of OMV and host yeast cells on the same side of the membrane strongly inhibited the growth of the cells (OD600 = 0.010±0.012 for 5.5 hr. incubation). These data suggest OMV activity against yeast cells depends on direct contact between the OMV and cells.

To assess the potential for contact-mediated transfer of antifungal activity, OMV binding to yeast cells was also studied using diIC18-labeled OMV. After incubation, OMV fluorescence was found associated with pellets of washed yeast cells in significant amounts (56±0.7% of total OMV fluorescence versus 7±4% in the absence of the yeast cells). Binding was also assessed by observation of S30 cells by fluorescence microscopy. Yeast cells, which were incubated with diIC18-labeled OMV and observed before washing, often showed a concentration of vesicles around the surface of the cells (Figure 8). Similarly, OMV labeled with an amine-reactive red fluorescent probe, also appeared to bind to some yeast cells that had been washed by sedimentation, although many fewer OMV remained attached to cells. (Figure 8).

**Figure 8:**
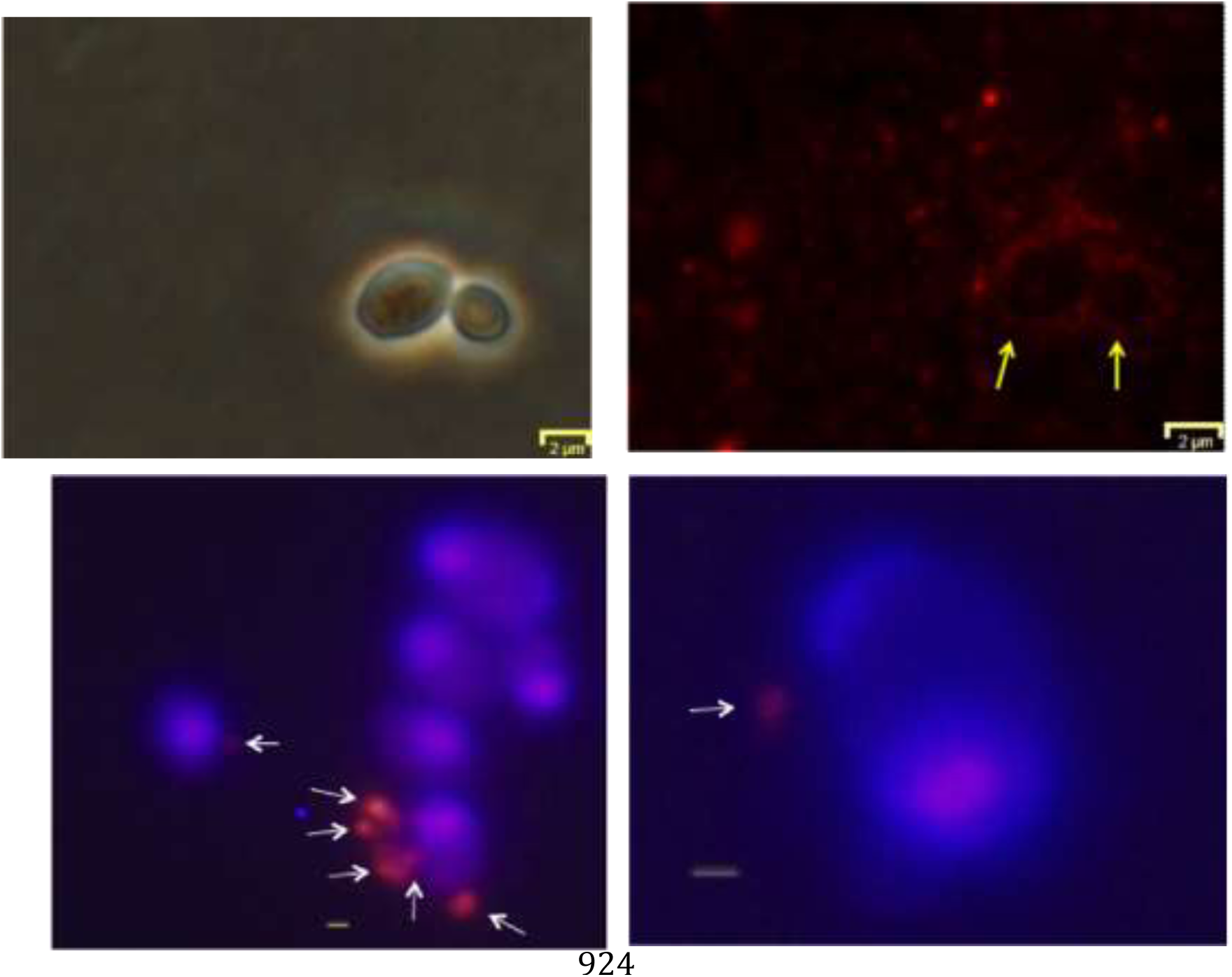
Binding of OMV to yeast cells. Top figures depict diIC18-labeled OMV (red fluorescence) interacting with yeast cells without washing. Top left and right figures are phase contrast and fluorescence versions of the same view. Yellow arrows indicate the position of two yeast cells. Bar represents 2 microns. Bottom photos: Hoechst 33342-labeled yeast cells (blue) were incubated with Alexa Fluor 555-NHS-labeled OMV (red) and then washed by centrifugation. Overlays of fluorescent images are shown. Grey bar represents 500 nm, white arrows indicate putative OMV or clusters of OMV bound to the cells.

Further studies with OMV labeled with membrane-localizing probes showed data consistent with transfer of these hydrophobic molecules to yeast cell membranes. When OMV labeled with the probes diIC18 or diOC18 were allowed to interact with the yeast cells for a period of time, and washed extensively, a dim diffuse fluorescence across the whole cell was observed. As noted above, a small fraction of these cells appeared to have bound, apparently intact OMV associated. However, essentially all the cells displayed the diffuse fluorescence, which was significantly above background as shown. It can therefore be proposed that this fluorescence may represent transfer of the OMV membrane-localized probes to membranes or other structures within the yeast cells. One interpretation of this result is that the OMV have some mechanism by which to transfer molecules from their membranes through the cell wall of the yeast cells and into the plasma membrane. Further investigation will be necessary to establish this paradigm.

### Identification of candidate molecules for antifungal activity

In an attempt to search for previously identified antifungal species, extracts were partially purified by thin layer chromatography and subjected to analysis by mass spectrometry. When C3 OMV were extracted with ethyl acetate, the concentrated extracts in methanol revealed a complex mixture of molecules with no apparently dominant species. Further purification was possible by a micro-preparative thin layer chromatography procedure. A UV-positive band was extracted from the plates and demonstrated to have yeast growth inhibitory activity. Analysis of an extracted band by mass spectrometry gave the data shown in Figure 10.

**Figure 9:**
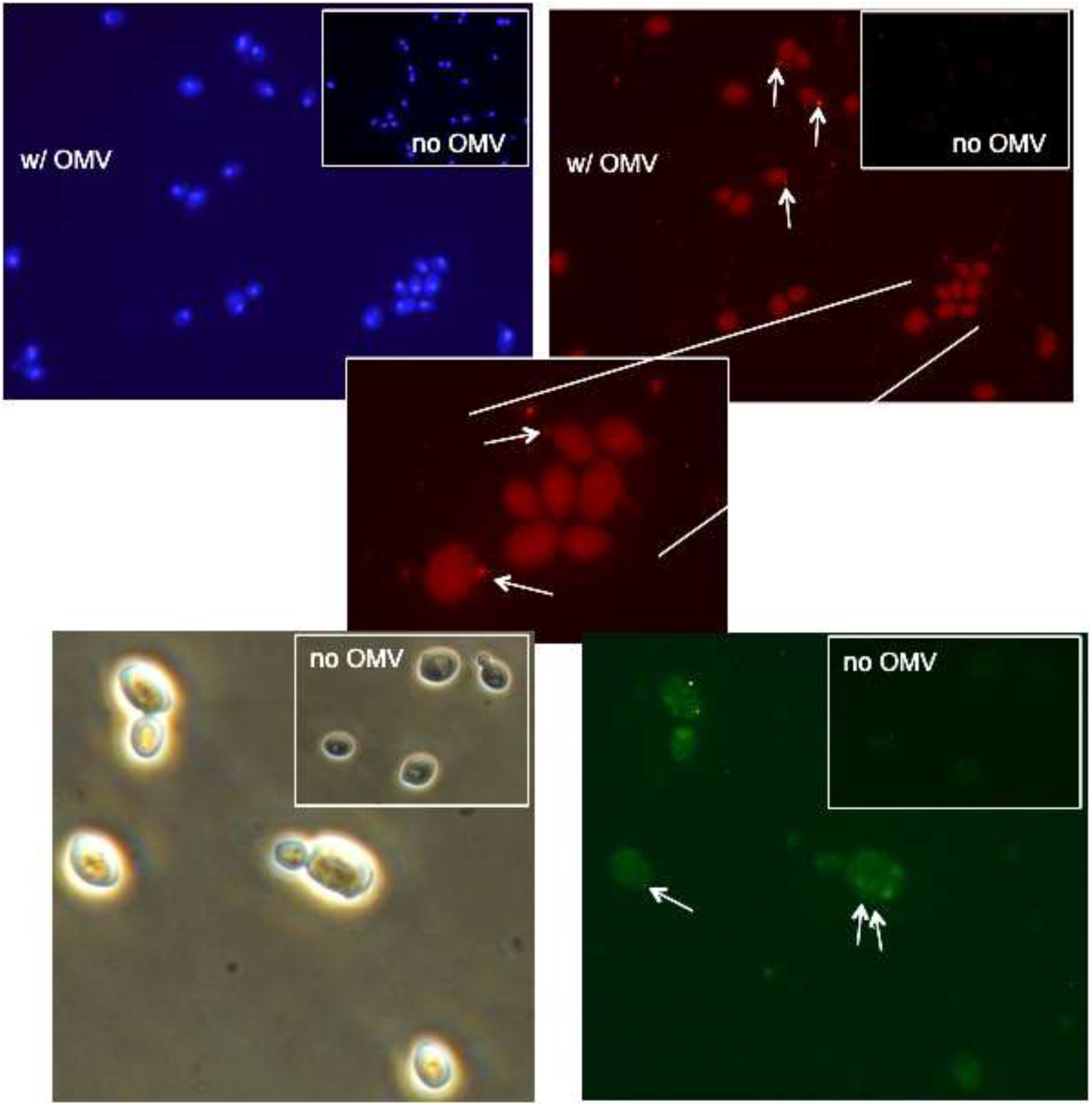
Apparent transfer of hydrophobic labels from C3 OMV to yeast cells. Yeast cells were labeled with Hoechst 33342 (upper photos, blue) or unlabeled (lower photos). OMV were labeled with diIC18 (red fluorescence, upper photos) or diOC18 (green fluorescence, lower photos) and allowed to interact with yeast cells. After washing to remove unbound OMV, yeast cells were observed by fluorescence or phase contrast microscopy. Insets in the upper right of each photo are shown to compare background fluorescence of samples with no added fluorescently labeled OMV. An expanded portion of one of the upper photos is shown in the middle panel. White arrows in the photos point to putative OMV bound to yeast cells. All fluorescent photos with and without OMV are compared under identical photomicrographic parameters and identical image enhancement parameters (brightness and contrast).

**Figure 10:**
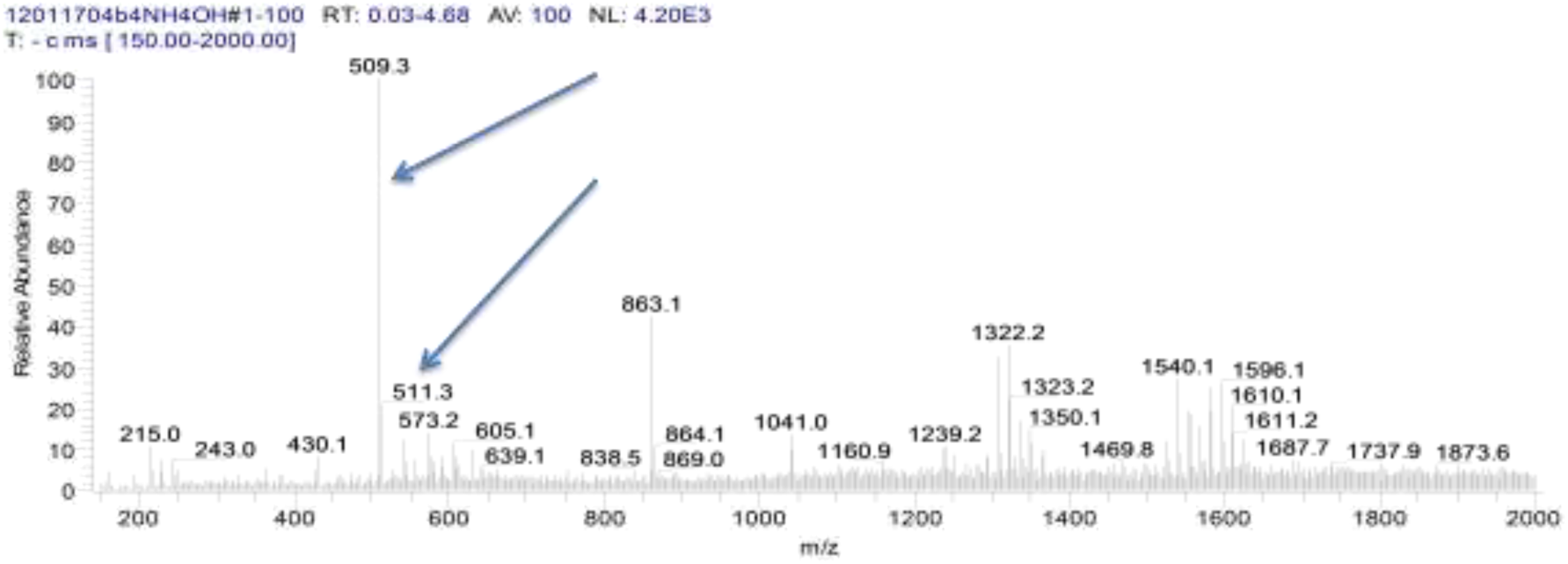
Negative-ion m/z spectrum of TLC-purified antifungal activity from C3 OMV. A UV positive band was obtained from a TLC separation of an OMV extract. The extracted band was dissolved in methanol for analysis by electrospray ionization mass spectroscopy as shown above. Arrows highlight peaks at 509.3 and 511.3 m/z (discussed in text). The peak at 863.1 m/z is an artifact that also appears in pure solvent spectra.

Because the TLC purification was performed in a strongly basic solvent, it was expected that negative ion products might be prominent in the analysis. In the negative ion mode, a prominent peak was observed at 509 m/z possibly consistent with alteramide B with a smaller peak at 511 possibly consistent with dihydromaltophilin (arrows, see discussion). A series of peaks is also seen in the range of 1332 and 1596 m/z. These groups of peaks appear to have multiplets separated by mass units of 14, the mass of CH_2_ units.

In the positive ion scans, some of the same peaks can be observed but apparently as disodium forms of the negative ions. For instance, peaks at 555 and 558 may correspond to peaks at 509 and 512 in the negative ion scans (data not shown). The potential identities of these compounds are discussed further below.

## Discussion

The existence of extracellular bacterial vesicles has been known for some time (8). But the identification of OMV as a potential part of the antifungal biocontrol activity of C3 fits a body of recent research elucidating the importance of OMV production as an extracellular mechanism of delivery of active molecules to host cells (12-24). Therefore, a role for OMV in the action of *Lysobacter* species fits an emerging paradigm in the field. The OMV identified here have some expected characteristics. For instance the ratio of LPS/protein in these preparations is particularly telling. The value observed for C3 (0.11 g/g) falls within a calculated range based on geometry and size (see Methods), and is also consistent with other measurements of this ratio in other OMV. For instance OMV from *Neisseria* and *Vibrio* have been reported to have a ratio of 0.06-0.08 and 0.08-0.09 (29,30), respectively. These relatively low ratios suggest that large amounts of protein are encapsulated in the interior or located on the surface of OMV. The apparent encapsulation of a substantial amount of chitinase activity in the interior of C3 OMV is consistent with this speculation.

OMV can act as long distance mediators of antifungal activity that would allow the bacterial cells to avoid fungal defenses while exerting a toxic effect on the target. However, the ability of C3 OMV to reach and deliver associated molecules to fungal cells brings up other questions. Specifically, the cell wall of fungi may be expected to inhibit the contact and transfer of molecules from vesicles. It cannot be fully ruled out that there is a mechanism by which the OMV can disrupt the cell wall. Our data with proteinase K digestion of OMV suggests that at least some portion of chitinase activity is located on the outside of the OMV, perhaps on the surface where it could mediate cell wall digestion. However, heat treatment, which destroyed chitinase activity and may do the same for other enzymes, did not affect the ability of OMV to deliver antibiotic activity under the conditions of these experiments. Although enzymatic activity may still prove to play a role, other mechanisms may need to be explored. Vesicular traffic across the cell wall of some species of fungi has been reported (31,32). Therefore, actual movement of OMV through the cell wall could be possible, though no evidence of this exists in the particular case of the experiments presented here.

Furthermore, it is not clear if the observation of antifungal activity in OMV under these conditions actually reflects the situation in a natural environment. On one hand, the observation of OMV under nutrient stress would seem to be consistent with conditions expected in natural environments, where nutrient broths do not exist. However, it is unclear when OMV would be released and how they might be directed to host cells, unless there is a mechanism of preferential binding. Previous data with other species have suggested that OMV can prime host cells for bacterial binding, which could apply in this case (33). Furthermore, it cannot be ruled out at this point that the observed production of OMV is really the consequence of a different mechanism of attack of the host cell that simply manifests as vesiculation of *Lysobacter* cells in broth culture. For instance, outer membrane bulging and protrusion may be a normal event when C3 cells make contact with host fungi without going all the way to vesicle production.

Though the mass spectrometry analysis alone does not positively identify any particular active molecule, the results strongly suggest two particular molecules. Alteramide B is the only C3 antifungal product specifically shown to possess significant activity against *S. cerevisiae* in previous work (6). The observed negative ion m/z of 509 is consistent with a single proton extraction from alteramide B. Therefore it is a likely candidate as an active molecule in these OMV. A small peak was also observed in the negative ion scan at 511 m/z. This species could be dihydromaltophilin, which has a molar mass of 512 in its unionized form. Finally a comment can be made about a series of peaks seen in the range of 1332 and 1596 m/z. These groups of peaks appear to have multiplets separated by mass units of 14, the expected mass of a CH_2_ unit. These m/z ratios could represent the existence of dimer ion clusters of phospholipids with a single negative charge, as has been observed previously (34). Because lipopolysaccharides are known to contain acyl groups of varying lengths, these multiplets may also represent lipid A species. Interestingly, the overall mass of these species could be consistent with partially deacylated lipid A based on comparison to the lipid A mass in other gram-negative bacteria (35). Deacylation of lipid A and LPS has been proposed to constitute part of a mechanism of OMV biogenesis in other bacterial species (35).

The fact that C3 OMV carries these small molecule antibiotics is a particularly important result. Though C3 antifungal antibiotics were previously thought to exist in the form of a protein-containing complex, identification of the relevant complexing entity as vesicular in nature is novel. The result has consequences in terms of the ongoing efforts to develop these antibiotics to treat plant or animal fungal diseases. At one level, the ability to concentrate and sequester the antibiotic activity of interest by a simple filtration process may help to speed the efforts to scale up and purify the relevant molecules. Additionally the very fact that the antifungal molecules are associated with OMV suggests that strategies to increase OMV production may also enhance antibiotic yields. On the other hand, the fact that the HSAF^-^ mutant still produces OMV, suggests that OMV and antibiotic production may not be tightly coupled. Further investigation would be needed to understand this point. In addition, the actual means of association of the antibiotic molecules with OMV membranes requires more study. One key parameter to understand is whether these antibiotics associate directly with the lipid bilayer of OMV as suggested for quorum signaling molecules (13) or with a membrane-associated protein. This localization may point the way to the best formulation strategies to ultimately accompany therapeutic development.

## MATERIALS AND METHODS

### Growth of bacteria and vesicle isolation

OMV were prepared from cell-free culture supernatants of the C3 strain of *Lysobacter enymogenes* (donated by the laboratory of Dr. Donald Kobayashi, Rutgers University) or in some cases from the K19 mutant strain designated HSAF^-^ (3) (kindly provided by the laboratory of Dr. Gary Yuen, University of Nebraska) in which the polyketide synthase of the HSAF polyketide pathway has been deleted by mutation. OMV were obtained from one of two media, 10% tryptic soy broth or minimal medium (MM). The latter is based on the medium used previously to obtain OMV from *Lysobacter sp.* XL1 (23). In some cases, MM was supplemented with 1% colloidal chitin. After overnight growth in one of these media, bacterial cells were removed by centrifugation of at 20,000 × g for 20 minutes (23). The remaining supernatants were either used directly for solvent extractions (see below) or processed to obtain OMV as follows.

OMV preparation from cell-free supernatants were performed either by direct ultracentrifugation at 29,000 rpm for 2 hour (Beckman SW40Ti swinging bucket rotor; ~140,000 × g, t/k-factor ≈ 0.88 min-rpm^2^), or the supernatants were first ultra-filtered through a 300 kDa cutoff polysulfone membrane (EMD Millipore, Billerica, MA) in a 400 ml stirred cell (Amicon model 8400). Both methods generated a small volume of solid material that was typically resuspended in Tris-buffered saline (TBS) and filtered through a 0.45 micron filter to remove any residual whole cells. Resuspension was followed by a smaller volume ultracentrifugation (Beckman AirFuge, A95 rotor, ~140,000 × g, 15 minutes; t/k-factor ≈ 1.25 min-rpm^2^) to obtain a concentrated pellet of OMV. Preparations were then washed by multiple centrifugation and resuspension steps (usually 2-3x). OMV yield was measured by protein content, typically by Bradford assay using bovine serum albumin as a standard. A typical yield from an overnight 300 ml culture in 10% TSB was approximately 0.5-0.7 mg of total OMV protein.

OMV were labeled when necessary using one of three different probes –Alexa Fluor 555 N-hydoxysuccinimide (Life Technologies, Grand Island, NY) to label amino groups and two hydrophobic probes, the green fluorescent dioctadecyloxacarbocyanine perchlorate (DiOC18) or red fluorescent *1,1^′^-*dioctadecyl-3,3,3′,3′-tetramethylindocarbocyanine perchlorate (DiIC18). DiOC18 in dimethylformamide was added to a vortexed solution of OMV at a ratio of 21 ng per µg of OMV protein (as measured by Bradford assay). Labeled OMV were pelleted and washed after labeling. OMV were labeled with DiIC18 in essentially the same way except that an ethanol solution was used. Alexa Fluor 555 NHS (~2 μg) was incubated with OMV (~50 μg protein) in sodium bicarbonate solution for 1 hr., followed by washing and resuspension in TBS. Fluorescence measurements were performed in 96 well plates using an Applied Biosystems Cytofuor series 4000 multi-well plate reader.

Samples for negative stain electron microscopy were prepared by resuspension of a high speed OMV pellet into a buffer of sodium cacodylate containing 2.5% glutaraldehyde for fixation and washed, followed by staining with uranyl formate. Stained samples were applied to glow discharged carbon/formvar-coated 300 mesh copper grids (EM Sciences, Hatfield, PA). Samples were observed using a JEM-100CXII electron microscope (JEOL) with the assistance of the Electron Microscopy Facility, Nelson Biology Laboratory, Rutgers University, New Brunswick, USA.

OMV particle size was measured using a *Zetasizer* Nano-ZS particle analyzer (Malvern Instruments, Westborough, MA with the assistance of Zoltan Szekely and Dan Myers). OMV samples were diluted approximately 10-100 fold for analyses, e.g. to approximately 10-20 μg/ml of protein in TBS.

#### LPS assay

Assay of LPS content was performed using a chromogenic version (Toxinsensor™) of the *Limulus* amebocyte lysate assay (Genscript, Piscataway, NJ). Standard curves based on *E. coli* LPS were generated to estimate the amount of LPS in C3 OMV samples.

#### Chitinase activity

Activity of OMV, from cells grown in MM with 1% chitin, was measured using a fluorogenic substrate, of 4-methylumbelliferyl β-D-N, N’, N”-triacetylchitotrioside (TC) (Sigma Aldrich, St. Louis, MO). Fluorescence is generated by enzymatic cleavage of the substrate and was measured in a 96 well plate format on an Applied Biosystems Cytofluor 4000 plate reader with excitation of 350 nm and emission of 460 nm. The relative fluorescence at defined time points within the linear range of fluorescence development was used to compare the activity of samples. For heat treatment experiments, OMV samples were incubated at 70 °C for 30 minutes before assaying for chitinase activity. Proteinase K (Sigma-Aldrich, St. Louis, MO) was used to assess accessibility of chitinase activity associated with OMV by comparison of incubation 48 hr. with proteinase K (~5 mg/ml) at 30 °C versus without. For freeze-thaw experiments, OMV samples were frozen in liquid nitrogen for at least 20 seconds and then thawed on a warm block in a process repeated ten times. OMV were washed free of soluble material after this treatment by sedimentation two times.

#### Sodium dodecyl sulfate polyacrylamide gel electrophoresis (SDS-PAGE)

Samples were prepared by dilution 1:1 with a 2x Laemmli sample buffer containing β-mercaptoethanol. Electrophoresis was performed on a Phast gel system (GE Healthcare, Life Sciences, Pittsburgh, PA) using an 8-25% polyacrylamide precast gel and SDS-containing buffer strips. Protein bands were visualized by silver staining. Western blotting was performed using the Phast Gel apparatus and a probe antibody against a peptide sequence (CTYTGGDKESQIPSS) in the pilA protein of C3 (Biomatik, Wilmington, DE, donated by Donald Kobayashi, Rutgers University). Detection was via chemiluminescence resulting from horseradish peroxidadase-coupled second antibody and a luminescent substrate (Genscript Lumisensor, Piscataway, NJ).

#### DNA assay

DNA content in OMV preparations was assessed either by standard extraction and precipitation methods involving ethanol or isopropanol, or by use of a fluorogenic detection method based on dye-binding to DNA. In the former, DNA was precipitated, resuspended in Tris-EDTA buffer and content was detected by optical density at 280 nm using a Nanodrop spectrophotometer. The identity of DNA was confirmed by comparing yields of OMV nucleic acid after incubation with DNase I or RNase A. In the fluorogenic assays, samples were incubated with Diamond™ Nucleic Acid Dye (Promega, Madison, WI), which can cross cell membranes. OMV were incubated with or without DNase I and then DNA content was measured by fluorescence (ex/em 485/580 nm) to assess protected or encapsulated DNA.

### Yeast Growth Assay

To measure yeast growth-inhibitory activity of OMV samples, the turbidity of *Saccharomyces cerevisiae* S30 cell cultures were measured. In initial experiments, approximately 10 ul each of OMVs and overnight S30 culture were incubated for 30 minutes at 30°C. The cultures were then expanded with 200μL YPD medium and incubated for 5.5 hours or overnight at 30°C. The optical density of 100 ul of each culture was diluted to 1 ml of medium was measured at 600nm (OD 600). In some cases the assay was modified by direct mixing of OMV and yeast into 200 μL of YPD medium followed by an overnight incubation. Both protocols gave similar results in terms of OMV activity in most cases. OMV from cells grown in 10% TSB showed similar specific activity to those from cells grown in MM with 1% chitin. For experiments to test contact dependence of OMV effects, small filtration devices (<1 cm diffusion length) with 100 kDa cutoff filters (Schleicher and Scheull, Centrex) were used and sterilely filled on opposite sides of the filter with solutions as indicated. The chamber with OMV alone or buffer alone contained 100 ul of YPD or Tris buffered saline with 10 mM glucose. The chamber with yeast cells contained 200 ul of YPD medium. Devices were incubated horizontally with shaking to facilitate liquid contact on both sides of the membrane. After 5.5 hours or overnight, 100 ul of the yeast-containing solution was diluted into 1 ml of YPD and OD 600 was measured. For experiments in which organic solvent extracts were being tested, ethanol-dissolved samples were used. The yeast cells were able to tolerate up to 15% (v/v) ethanol in YPD medium.

### Fusarium growth assay

*Fusarium subglutinans* (kindly provided by Marshall Bergen and James White, Rutgers University) was initially cultured in YPD liquid medium and then spread onto YPD agar plates. A 4 mm plug from the plate was excised and laid upside down onto the surface of a fresh YPD plate so that hyphae grew from the center of the plate. After sufficient growth, wells were created approximately 5 mm from the edge of the growing mycelia, and samples (50 μL) were loaded into the wells for testing. Growth in the region of the wells was assessed by visual observation.

### OMV binding to yeast cells

OMV were labeled for yeast binding assays with an ethanolic solution of DiIC18, as described above, at a ratio of 20 ng per µg of OMV protein with a final ethanol concentration of 0.1% (v/v). This mixture was incubated at 37 C for 1 hr. Labeled OMV were pelleted (140,000 × g) and resuspended in 400 µl TBS. The S30 strain of the yeast *Saccharomyces cerevisiae* was grown overnight in YPD medium, collected and washed by sedimentation and resuspended in TBS with 10 mM glucose. Typically 10 µl of OMV was mixed with 20 µl of yeast cells diluted into 1 ml of TBS with glucose. Association with yeast was measured by incubation at 30 °C followed by collection of yeast cells by centrifugation at 4500 × g. Fluorescence of the DiIC18 associated with the yeast pellet was used as a measure of association.

For OMV binding experiments by microscopy, the S30 strain of the yeast *Saccharomyces cerevisiae* was grown overnight in YPD medium and collected and washed by sedimentation. 10 μl of the washed overnight culture was incubated with 10 μl of labeled OMV preparations (approximately 50 ng total protein) for 2 hours, diluted into 1 ml of Tris buffered saline (TBS) with 10 mM glucose and the yeast cells collected by centrifugation at 1000 × g. After resuspension in 10 μl of TBS/glucose, the samples were used for fluorescence microscopy. In some cases, samples were observed without dilution and removal of unbound OMV. Samples were observed using an Olympus FSX100 inverted epifluorescence microscope.

### Solvent extractions of antifungal activity

Bligh-Dyer extractions to analyze lipids were performed as described (27) with appropriate volume adjustments to accommodate sample size.

Ethyl acetate was used to extract preparations at various stages of purification with variations on previously published methods (28). For the large volume cell-free supernatants used in analysis of activity in Figure 3, 125 ml of ultrafiltrate (see above) or a resuspension of the material on the filter surface into 125 ml of 10% TSB was used for extraction. Medium was extracted 3 times with 50 ml of ethyl acetate using a separatory funnel. The collected upper phase of solvent was evaporated and the entire extraction was redissolved into 250 μL of ethanol for activity testing.

Some washed OMV preparations were extracted with ethyl acetate in a smaller volume format. Preparations were diluted to approximately 100 μg/ml of protein before extraction into an equal volume of ethyl acetate. After removal of solvent, the solids were redissolved into ethanol for activity assays or methanol for mass spectrometry or further purification by thin layer chromatography.

#### Thin layer chromatography

Samples obtained from Bligh Dyer extractions to test for OMV phospholipid content (data in Figure 3) were concentrated and spotted onto silica gel plates (Analtech Uniplate 250 micron HF; Sigma Aldrich, St. Louis, MO) and run in the solvent system 65:25:2.5 chloroform: methanol:ammonium hydroxide. Phospholipid standards were 1-palmitoyl-2-oleoyl-*sn*-glycero-3-phosphocholine (POPC) and 1-palmitoyl-2-oleoyl-*sn*-glycero-3-phospho-(1′-*rac*-glycerol) (POPG) from Avanti Polar Lipids (Birmingham, AL).

For purification of active antifungal molecules by ethyl acetate extraction and analysis by mass spectrometry, the solvent system 65:25:5 chloroform:methanol:ammonium hydroxide was initially used. On preparative scale plates (Analtech Uniplate 20 × 20 2000 micron silica gel GF with UV 254 nm fluorescence), this system yielded a UV-positive band at approximately Rf 0.11 which could be extracted into methanol, redissolved in ethanol and demonstrated to possess activity against yeast cells. Subsequently for MS analysis, samples were purified using a solvent system of 65:30:5 chloroform:methanol:20% ammonium hydroxide, which resulted in an Rf of 0.55 for the active compound (MS grade HPTLC silica gel 60 F_254_; EMD Merck, Burlington, MA). This band was extracted into methanol for analysis by mass spectrometry.

Mass spectrometry was performed on a Finningan LCQ Duo electrospray ionization instrument (Department of Chemistry, Rutgers University; thanks to Alexei Ermakov for assistance).

### Calculations

OMV LPS/protein calculation:

An estimation of range of possible ratios for LPS/protein was calculated based on assumption of spherical vesicles of about 135 nm in diameter with an outer monolayer consisting of LPS and 50% by weight of protein in the membrane. Calculation of maximum possible value: The maximum LPS/protein value would be in empty vesicles. If the membrane is assumed to be 50% by weight protein, then, of the remaining 50%, more than half is LPS. We assume 33% of total weight is LPS (because of the large LPS molecular weight). So maximum LPS/protein g/g is 0.66.

The lowest possible LPS/protein ratio would come from OMV packed with a maximum amount of protein in the interior of the vesicle. An internal radius of 64 nm is assumed with 3 nm membrane thickness. Then the internal volume = 4/3* 3.14* (64 nm)^3^ = 1.1e6 nm^3^. An estimation of the protein in the membrane can be made from the total membrane volume 4/3* 3.14 (67 nm)^3^ - 4/3* 3.14 (64 nm)^3^ = 162,000 nm^3^. So the ratio of protein in the filled vesicle to that in the empty vesicle would be (1.1+0.16)/0.16 = 7.9. By this estimate the minimum LPS/protein ratio would be 0.66/7.9 = 0.08. These are clearly very rough estimates, but finding an LPS/protein ratio in this range would give a modicum of confidence in the measurement.

## Acknowledgements

This work was funded by the Byrne Seminar program, Department of Plant Biology, Rutgers Master in Business and Science program, and personal contributions from Paul Meers. Equipment and supplies were donated by Transave, Inc., Anthony Scotto, Ph.D. and Barbara A. Zilinskas, Ph.D. Students who contributed to this work but could not be reached for authorship approval are Shanee Grant and David Itenberg.

